# Analysis of M4 transmembrane domains in NMDA receptor function: a negative allosteric modulation site at the GluN1 M4 is determining the efficiency of neurosteroid modulation

**DOI:** 10.1101/2021.03.15.435138

**Authors:** Kai Langer, Adriana Müller-Längle, Jannik Wempe, Bodo Laube

## Abstract

Ionotropic glutamate receptors (iGluRs) are tetrameric ligand-gated ion channels that play a crucial role in excitatory synaptic transmission in the central nervous system. Each subunit contributes with three transmembrane domains (M1, M3, and M4) and a pore loop (M2) forming the channel pore. Recent studies suggest that the architecture of all eukaryotic iGluRs derives from a common prokaryotic ancestral receptor that lacks M4 and consists only of the transmembrane domain segments M1-M3. Although a crucial contribution of M4 to the assembly and trafficking of iGluRs is suspected, the role of this additionally evolved domain in receptor function remains controversial. Here, we investigated how deletions and mutations of M4 in members of the NMDA receptor subfamily, the conventional heteromeric GluN1/GluN2 and glycine-gated GluN1/GluN3 NMDA receptors, affect expression and function in *Xenopus* oocytes. We show that deletion of M4 in the GluN1, GluN2, or GluN3 subunit, despite retained receptor assembly and cell surface expression, results in nonfunctional membrane receptors. Coexpression of the corresponding M4 domains as an isolated peptide in M4-deleted receptors rescued receptor function of GluN1/GluN2A NMDARs without altering the affinity of glutamate or glycine. Substitution of non-conserved residues and insertion of interhelical disulfide bridges confirmed the proximity of positions M813 and F817 in M4 of GluN1 to residues of the TMs of neighboring subunits. Electrophysiological analyses of agonist-induced receptor function and its modulation by the neurosteroid pregnenolone sulfate (PS) at mutations of the GluN1-M4/GluN2/3-TM interface indicate a crucial role of interdomain interactions in the functional coupling of M4 to the nuclear receptor and the modulatory effect of PS. Our results show that although the M4 domains in NMDA receptors are not important for receptor assembly and surface expression, residues at the subunit interface are substantially involved in M4 recognition to the core receptor and regulation of PS efficacy. Because mutations in the M4 of GluN1 specifically resulted in loss of PS-induced inhibition of NMDA receptor currents, our results point to distinct roles of M4s in NMDA receptor modulation and highlight the importance of evolutionarily newly evolved M4s for selective *in vivo* modulation of glutamate- and glycine-activated NMDA receptors by steroids.

**Highlights:**

- The role of the M4 transmembrane domain in the assembly and function of ionotropic glutamate receptors remains controversial
- Here we show that deletion of M4 in glutamate-gated GluN1/GluN2A and glycine-gated GluN1/GluN3A receptors results in nonfunctional NMDA receptors with retained surface expression
- The functional loss in M4-deleted GluN1/GluN2A receptors is rescued without affecting agonist affinity by a M4 transmembrane domain of the respective subunit expressed as an isolated peptide
- Specific interactions in the M4 interfaces with the M1 and M3 domain of the adjacent subunit are required for the recognition of the isolated M4 and the functional rescue
- Finally, the M4 domain-interfaces of GluN1 determine the negative modulatory effect of pregnenolone sulfate in glutamate-gated GluN1/GluN2A and glycine-gated GluN1/GluN3A NMDA receptors

## 1. Introduction

The majority of excitatory activity in the central nervous system (CNS) is mediated by the neurotransmitter glutamate. At postsynaptic neurons, glutamate binds to a wide range of glutamate receptors. Ionotropic glutamate receptors (iGluRs) are one of the major classes of cation-selective ion channels and are divided into four main classes, the AMPA receptors (GluA1-A4), kainate receptors (GluK1-K5), NMDA receptors (GluN1, GluN2A-D, and GluN3A-B), and the delta receptors (GluD1-D2). These receptors are widely distributed in the CNS and play important roles in CNS development, the formation of respiratory and locomotor rhythms, and processes such as learning, memory, and neuroplasticity (Collingridge and Bliss, 1995; Yashiro and Philpot, 2008). iGluRs are characterized by a tetrameric structure of four identical or similar subunits that all follow the same scheme. The subunits are modular and consist of an extracellular amino-terminal domain (ATD), a ligand-binding domain (LBD), the transmembrane domain (TMD), and an intracellular carboxy-terminal domain (CTD). The transmembrane domain consists mainly of the secondary structure motif of α-helices that embed into the cell membrane, anchoring the receptor in the synaptic membrane (Sheng and Pak, 1999). The TMD of iGluRs consists of transmembrane helices M1, M3, and M4 and a membrane loop named M2. The cation-selective pore in the center of the assembled receptor is formed by helices M1, M2, and M3 (Traynelis and Wollmuth L.P., McBain C.J., Menniti F.S., Vance K.M., Ogden K.K., Hansen K.B., Yuan H., Myers J.M., 2010). With this pore structure, they inversely resemble bacterial potassium channels, assuming an evolutionary connection (Wo and Oswald, 1995). Thus, it has been shown that it is possible to generate functional chimeras between iGluRs and potassium channels lacking the M4 domain (Schönrock et al., 2019). These findings are also consistent with the functional subunit structure of a bacterial iGluR (i.e., GluR0 from *Synechocystis*) consisting of subunits with only two transmembrane domains (M1 and M3) lacking M4.

While the function of helices M1 - M3 is well understood, the role of the evolutionarily newly formed M4 is still unclear. Structurally, it is distant from the other membrane helices of its own subunit and can only interact with M1 or M3 of the neighboring subunit. Hence, different hypotheses about the exact role of M4 are currently discussed. For AMPA receptors, there is conjecture about the involvement of the M4 domain in assembly according to the “dimer of dimers” principle. Subunits lacking the M4 domain caused receptors to stack in the endoplasmic reticulum and fail to tetramerize, whereas subunit dimerization appears to function correctly. Single substitutions in the M4 domain at interaction sites near the M1 and M3 segments of adjacent subunits impaired receptor association, suggesting a role for these positions within receptor assembly (Salussolia et al., 2013). For NMDA receptors, there is evidence for M4 involvement in masking retention signals in the M3 domain of the adjacent subunit within the endoplasmic reticulum, leading to a strong decrease in receptor surface expression in the absence of M4 (Horak et al., 2008). Consistent with this finding, removal of the M4 domain from GluN1/GluN2A receptors resulted in functionally undetectable receptor variants (Schorge and Colquhoun, 2003). Moreover, tryptophan screens of the M4 domain identified different positions at the extracellular end of the M4 domain that influenced GluN1/GluN2A receptor functionality, whereas positions at the more intracellular end of the M4 segment showed a higher impact on surface expression (Amin et al., 2017). In addition to being involved in NMDAR assembly and function, the M4 domain appears to be critical for receptor modulation. The neurosteroid pregnenolone sulfate has a bivalent function at NMDARs that is thought to be mediated by two distinct binding sites. First, at GluN2A or GluN2B subunit containing NMDARs PS acts as a positive allosteric modulator (PAM); second, at GluN2C or GluN2D subunit containing NMDARs PS acts as a negative allosteric modulator (NAM). The PAM binding site of PS is thought to be in the transmembrane region of the GluN2 subunit (Horak et al., 2006; Wilding et al., 2016). Further experiments showed that there are two obligatory regions for the effects of PS on all combinations of NMDA receptors. The J/K helix connecting the LBD to the M4 and the M4 domain itself are required for modulation (Jang et al., 2004). Recent findings through molecular dynamics simulations and alanine screening indicate that the binding site responsible for positive modulation of GluN1/GluN2A/B indeed appears to be localized at the GluN2 M1/M4 interface. These simulations also raise the prospect of another binding site at the appropriate positions for the GluN1 M1/M4 interface that does not carry the PAM effect (Krausova et al., 2020).

In this study, we addressed the question of the extent to which the M4 domain of NMDARs is required for receptor function. To answer this question, we used an approach of M4 truncation, separate M4 segment co-expression, and point mutations in NMDA receptors to gain insight into the influence of the M4 domain on functionality, assembly, and steroid modulation. To this end, we exploited the ability of separate M4 expression to rescue the functionality of the M4-truncated core receptor. Using this approach, we clearly demonstrated that the M4 domain is not involved in assembly and surface targeting but is essential for NMDA receptor functionality. Furthermore, we found two possible attachment sites of the GluN1 M4 to the M1 and M3 domains of the GluN2A that have a strong influence on the rescue effect. In addition, we identified a specific steroid recognition site at the GluN1 M4 domain that mediates the NAM effect of PS on NMDA receptors.

## 2. Material and Methods

Chemicals were purchased from Sigma (Taufkirchen, Germany). Restriction enzymes, Phusion polymerase, and T4 ligase were purchased from Thermo Fisher (Waltham, USA).

### 2.1 DNA constructs, oocyte expression and electrophysiology

The GluN1-1a (splice variant 1a), GluN2A, GluN2D, and GluN3A expression constructs in the pNKS2 vector used have been described previously (Honer et al., 1998; Madry et al., 2007b). The M4-deleted GluN1^1-802^ (named GluN1^ΔM4^) and GluN3A^1-893^ (GluN3A^ΔM4^) and the M4 constructs GluN1^802-925^ (named M4^N1^) and GluN3A^893-1092^ (M4^N3A^) tagged with the respective subunit signal peptide were generated using a Nhe I restriction site. GluN2A^1-803^ (GluN2A^ΔM4^) and GluN2A^804-1464^ (M4^N2A^) constructs were generated by overhang PCR. Biochemical expression analyses were performed using a C-terminal truncated GluN2A^1-929^ construct (GluN2A*) to avoid the confounding effect of the C-terminus during purification steps (Mesic et al., 2016). Point mutations were generated by missmatch PCR. All constructs were confirmed by DNA sequencing (Seqlab, Göttingen, Germany). After plasmid linearization with NotI, cRNA was synthesized using the Amplicap-Max™-SP6 High Yield Message Maker Kit from Cellscript (Madison, Wi, USA) as described by Mesic et al., (2016). Oocytes were obtained from female *Xenopus laevis* after anesthesia with 0.2% tricaine in water after the approval of the Technische Universtät Darmstadt (Agreement V54-19c20/15 DA8/No. 20). Oocytes were isolated and preserved as previously described (Laube et al., 1997). For electrophysiological analysis, oocytes were injected with 50 ng in a volume of 50 nl of the respective wt, M4-deleted, and/or M4 cRNAs in a 1:1:1 ratio. Whole-cell currents were recorded 3 days after injection by two-electrode voltage clamp (TEVC) according to Laube et al (1997). In brief, TEVC measurements were performed at room temperature using a GeneClamp 500B amplifier and a Digidata 1322A as an A-D converter. Measurements were recorded using Clampex 10.7 (Molecular Devices, San Jose, USA) at 5 kHz after low-pass filtering at 200 Hz. Microelectrodes with a resistance of 0.8-2.3 MΩ were filled with 3 M KCl. Oocytes were clamped at −70 mV. For application, compounds were dissolved in Ringer’s solution, except for pregnenolone sulfate, whose stock solution (100 mM) was prepared in DMSO. For I_max_ current determination of GluN1/GluN2A receptors, glycine and glutamate (100μM each) were coapplied. For dose-response analysis, the respective coagonist was applied at 100 μM. Measurements of GluN1/GluN3A receptors were made with 10mM glycine after preapplication of MDL-29951 (Madry et al., 2007a). For dithiothreitol (DTT) treatments, oocytes were superfused with 2 mM DTT for 100 s before agonist application in the presence of 2 mM DTT, as described by Lynagh et al (2013). Pregnenolone sulfate (PS) was always preapplied 50 s before application of the respective agonists. TEVC recordings were analyzed using Clampfit 10.7 (Axon industries). For dose-response analysis, currents were normalized to the maximum inducible current (I_max_) and fitted with variable slope nonlinear regression in GraphPad Prism 7.0 (GraphPad Software Inc., La Jolla, USA) as previously described (Lynagh and Laube, 2014).

### 2.2 Labeling, purification and SDS-PAGE of NMDA receptor complexes

For expression analysis, surface receptors were labeled with Pierce^™^ Premium Grade Sulfo NHS-SS-Biotin (Thermofisher, Waltham, USA) and purified using Streptavidin High Performance Spintrap^™^ (Sigma-Aldrich, St.Louis, USA). If samples were also treated with DTT, they were incubated with 100 mM DTT (Stocksolution 2mM DTT in ddH_2_O) for 20 min at 56 °C. Isolated surface proteins were separated on linear 10% SDS-PAGE gels. PVDF membranes and the Trans-Blot^®^Turbo^™^Blotting System (Biorad, Hercules, USA) were used for Western blot analysis. Two different antibodies were used as primary antibodies, firstly the GluN1-CTD was addressed with an antibody from Merck Millipore (Darmstadt, Germany) diluted 1:1000 in TBS, and secondly the GluN1-NTD epitope was addressed with an antibody from Alomone Labs (Jerusalem, Israel) 1:500 in TBS. A horseradish peroxidase-labeled secondary antibody (1:20000 in TBS) detecting mouse or rabbit IgG was used. Immunoreactive bands were visualized with the Pierce^™^ ECL Western Blotting Substrate (Thermofisher, Waltham, USA) using the ChemiDoc MP Imaging System (Biorad, Hercules, USA). Metabolic labeling with [35S]methionine (0.2Mbq per oocyte; >40 TBq/mmol, Amersham Biosciences) was performed as previously described (Schüler et al., 2008). Purification of C-terminal His6-tag-labeled GluN1 and GluN1^ΔM4^ by Ni^2+^-NTA agarose (Qiagen) chromatography was performed as in (Madry et al., 2007a). [35S]-Methionine-labeled protein samples were solubilized in SDS sample buffer containing 20 mM dithiothreitol and electrophoresed in parallel with molecular mass markers (SeeBlue^®^ Plus2 Pre-Stained Standard, Invitrogen) on 8% Tricine-SDS polyacrylamide gels. Gels were blotted, fixed, dried, and exposed to BioMax MR films (Kodak, Stuttgart, Germany) at −80 °C. The radioactivity of each protein band was quantified using a PhosphorImager (Molecular Dynamics) and analyzed using the ImageQuant software package. Cy5-NHS labeling (Amersham Biosciences) and subsequent SDS-PAGE were performed as described in (Mesic et al., 2016) and scanned with a gel imager (Typhoon 9400, Amersham Biosciences) as described (Madry et al., 2007a). To distinguish between mature and immature receptor complexes, 10 μl of affinity-purified receptor was incubated in reducing sample buffer (20 mM DTT, 1% (w/v) SDS) with 1% (w/v) octylglucoside containing 5 U endoglycosidase H (Endo H) or peptide: N-glycosidase F (PNGase F; both NEB, Frankfurt, Germany) at 37°C for 1 h, and protein samples were analyzed by SDS-PAGE as described above.

### 2.3 Sequence and structural analysis

Sequence alignments of the GluN1, GluN2, and GluN3 receptor sequences (taken from UniProt) were performed using the Multiple Sequence Alignment Tool from EMBL-EBI (Cambridge, UK). UCSF Chimera (Pettersen et al., 2004) was used for structural analysis, with the GluN1/GluN2A/GluN2B structure 5UP2 (Lü et al., 2017). The distances between the Cα-atoms of the amino acids were determined using the distance tool in Chimera. Images were generated using PyMOL 1.2 (http://www.pymol.org).

### 2.4 Statistical analysis

Statistical analysis was performed using GraphPad Prism 7.0 (GraphPad Software Inc., La Jolla, USA). A Gaussian distribution was assumed for the values obtained. Paired or unpaired Student’s t-test was used to determine significances. p < 0.05 (*), p < 0.01(**), p < 0.001 (***), and p <0.0001 (****). Values shown represent means ± SEM.

## 3. Results

### 3.1 The M4 transmembrane domain is essential for the function of glutamate-gated GluN1/GluN2A NMDA receptors

To analyze the function of M4 domains in NMDA receptors, we first examined the functional properties of the glycine-binding GluN1 subunit truncated by the M4 domain (GluN1^ΔM4^; for illustration, see Fig. 1A and MatMet) after coexpression with the glutamate-binding wild-type (wt) GluN2A subunit. In contrast to wt GluN1/GluN2A receptors (Fig. 1B), no currents could be measured in GluN1^ΔM4^/GluN2A subunit-expressing *Xenopus laevis* oocytes after application of saturating concentrations of glutamate and glycine by two-electrode voltage clamping (TEVC) (Fig. 1B). However, coexpression of GluN1^ΔM4^/GluN2A receptors in the presence of a protein fragment containing the M4 transmembrane domain of GluN1 (M4^N1^) resulted in agonist-induced currents (Fig. 1B). To investigate whether M4 protein fragments from other NMDAR subunits would also rescue GluN1^ΔM4^/GluN2A receptor function, we coexpressed homologous constructs of the M4 of GluN2A (M4^N2A^) and GluN3A (M4^N3A^) subunits. No detectable currents were obtained for either M4 fragment in GluN1^ΔM4^/GluN2A receptors after coexpression, indicating a specific role of the M4^N1^ domain in the rescue of GluN1^ΔM4^/GluN2A receptors. To analyze the pharmacological properties of GluN1^ΔM4^/GluN2A receptors in the presence of the M4^N1^ protein fragment, the maximum inducible currents (I_max_) and the respective EC50 values of agonists were measured in wt-GluN1/GluN2A and GluN1^ΔM4^+M4^N1^/GluN2A receptors. GluN1^ΔM4^+M4^N1^/GluN2A-expressing oocytes showed lower I_max_ than wt-GluN1/GluN2A receptors upon application of saturating concentrations of glutamate and glycine (GluN1/GluN2A: 2.81±0.4 μA, vs. GluN1^△M4^+M4^N1^/GluN2A: 0.15±0.04 μA, t(9)= 5.663; p<0.001; Fig. 1B). In contrast, EC50 values of glutamate and glycine affinities were not altered compared to wt GluN1/GluN2A receptor (glutamate: 2.45±0.3 μM vs. 2.3±0.3 μM; glycine: 1.8±0.25 μM vs. 2.35±0.2 μM for GluN1/GluN2A and GluN1^ΔM4^+M4^N1^/GluN2A receptors, respectively; Glu: t(5)= 0.447; p = 0.674 and Gly: t(7)= 1.584; p= 1.572 see Fig. 1C). In conclusion, only co-expression of the M4 protein fragment of the GluN1 subunit rescued GluN1^ΔM4^/GluN2A channel function without altering glutamate and glycine affinities.

**Figure 1:**
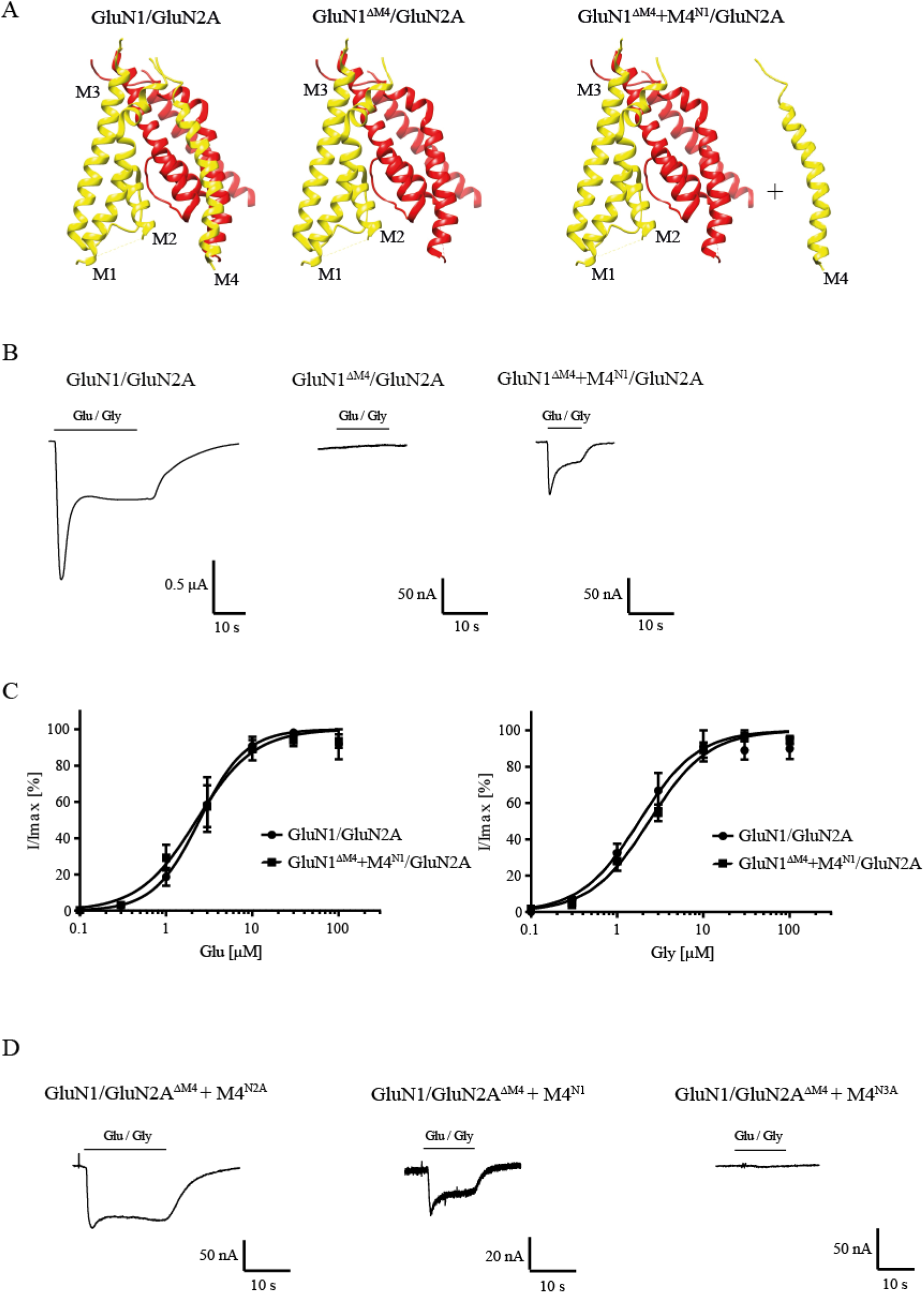
Functional characterization of M4-deleted and M4-co-expressed GluN1/GluN2A receptors. (A) Structure of the TMD region of a GluN1(yellow)/GluN2A(red) receptor showing the peripheral localization of the M4 domain compared to M1, M2 and M3 of its own subunit. Structure 2 and 3 show the M4-truncated receptor (GluN1^ΔM4^/GluN2A) and exemplary M4-truncated receptor with the separate M4 segment (GluN1^ΔM4^+M4^N1^/GluN2A). Structural analysis was performed from PDB 5UP2 using UCSF Chimera. (B) Representative TEVC images of the wt GluN1/GluN2A receptor, the non-functional truncated GluN1^ΔM4^/GluN2A, and the GluN1^ΔM4^+M4^N1^/GluN2A receptor rescued by M4 segment co-expression. GluN1^ΔM4^+M4^N1^/GluN2A showed similar kinetic properties compared with wt but a reduced I_max_ current. (C) Dose-response analysis of glutamate and glycine for wt and GluN1^ΔM4^+M4^N1^/GluN2A showed no significant differences in agonist affinity (EC_50_ glutamate: 2.45±0.3 μM (n = 3) vs. 2.3±0.2 μM (n = 4); EC_50_ glycine: 1.8±0.25 μM (n = 3) vs. 2.35±0.2 μM (n = 5) for GluN1/GluN2A and GluN1^ΔM4^+M4^N1^/GluN2A; p>0.05). (D) Representative TEVC images of GluN1/GluN2A^ΔM4^+M4^N2A^ and GluN1/GluN2A^ΔM4^+M4^N1^ show rescued functionality of GluN2A-M4 truncated receptors with M4^N1^ or M4^N2A^. No measurable currents could be obtained for GluN1/GluN2A^ΔM4^+M4^N3A^, highlighting the differences between the M4 domains of GluN1, GluN2A, and GluN3A. Data represent mean values ±SEM.

To analyze the role of the M4 of GluN2 subunits in GluN1/GluN2 receptor function, we examined the functional properties of the glutamate-binding GluN2A subunit (GluN2A^ΔM4^) truncated by the M4 domain after coexpression with the wt GluN1 subunit. Again, only the presence of the M4 protein fragment of GluN2A (M4^N2A^) resulted in inducible currents in GluN1/GluN2A^ΔM4^ receptor-expressing oocytes with an I_max_ of 0.24±0.1 μA (Fig 1D). In contrast to our results with the GluN1^△M4^ construct, where only the M4 of the GluN1 subunit rescued channel function, coexpression of the M4 of the GluN1 subunit (M4^N1^) also rescued receptor function of the GluN2A^ΔM4^ construct (I_max_ 0.016±0.007 μA, Fig 1D). However, coexpression of the M4^N3A^ fragment with GluN1/GluN2A^ΔM4^ receptors showed no rescue effect (Fig. 1D). Thus, both the M4 transmembrane domain of GluN1 and of GluN2A could rescue the channel function of GluN1/GluN2A^ΔM4^ receptors after coexpression. In summary, our analyses of the rescue of M4-truncated GluN1/GluN2A NMDA receptors by separately expressed M4 protein fragments indicate that i) specific recognition interactions must exist between the M4 and the nuclear receptor, since only in the presence of selected NMDAR-M4 fragments the deleted GluN1/GluN2A receptors were functional, and ii) the basic pharmacological properties were not altered, since the glutamate and glycine affinities remained the same compared with wt GluN1/GluN2A.

### 3.2 M4-deleted NMDAR subunits retain receptor surface expression

To investigate whether M4-truncated glycine-gated GluN1/GluN3 NMDARs are also rescued in function by M4 fragments, we analyzed our GluN1^ΔM4^ with wt GluN3A subunit in the absence and presence of M4^N1^. In contrast to our results with the M4-truncated GluN1/GluN2A receptors, neither application of the agonist glycine (1 mM) alone nor in combination with the potentiating ligand MDL-29951 (0.2 μM, see Madry et al., 2007a) caused detectable currents in GluN1^ΔM4^/GluN3A receptors in the presence of M4^N1^ (Fig. 2A). To analyze whether the GluN1^ΔM4^-assembles with the GluN3A subunit, we performed SDS-PAGE after metal affinity chromatography with a C-terminal hexahistidyl-tagged GluN3A subunit (GluN3A-His) after metabolic [35S]methionine labeling (see Material and Methods). Coexpression of GluN3A-His with either the GluN1 or the GluN1^ΔM4^ subunit resulted in two [35S]methionine-labeled subunits each with similar 1:1 intensities (Fig. 2B and data not shown) after autoradiographic analysis of the radioactive bands based on the total number of methionine residues per subunit (30 per GluN1, 25 per GluN1^ΔM4^, and 33 per GluN3A; see Mesic et al,. 2016). This indicated an unchanged assembly behavior of the GluN1^ΔM4^/GluN3A receptor. To understand the differential rescue effect of the M4^N1^ fragment in GluN1^ΔM4^/GluN2A and GluN1^ΔM4^/GluN3A receptor function, we examined the expression pattern of the GluN1^ΔM4^ and GluN3A-His subunits after coexpression with the M4^N1^ fragment (Fig. 2C). In contrast to the result with the GluN1^ΔM4^/GluN3A receptor in the absence of M4^N1^, surprisingly, both the GluN3A and GluN1^ΔM4^ subunits showed a marked reduction in total protein amount after coexpression with the M4^N1^ fragment (compare lanes 1 and 4 in Fig. 2C). We therefore performed the reverse experiment and analyzed the biochemical and functional properties of the M4-truncated GluN3A subunit (GluN3A^ΔM4^) after coexpression with the wt-GluN1 subunit in the absence and presence of the M4^N3A^ fragment. Similar to GluN1^ΔM4^/GluN3A receptors, coexpression of GluN1/GluN3A^ΔM4^ with the M4^N3A^ fragment resulted in nonfunctional channels and a comparable decrease in protein expression of GluN1/GluN3A^ΔM4^ receptors (see Fig. 2D compared with 2C). These data suggest that the respective M4 fragments impede protein expression of GluN1^ΔM4^/GluN3A and GluN1/GluN3A^ΔM4^ receptors, respectively.

**Figure 2:**
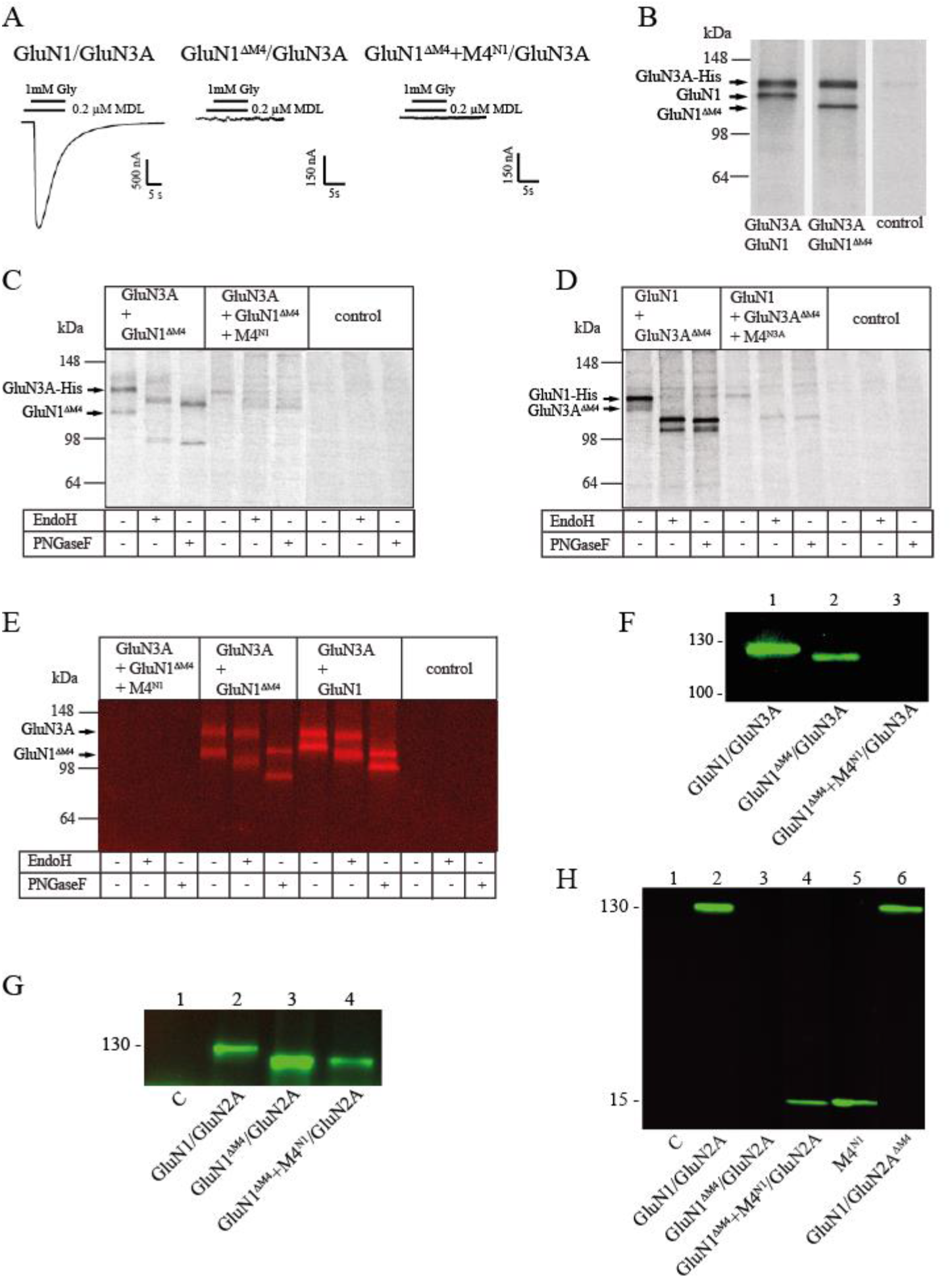
Effect of M4 domain on surface expression of NMDA receptors. (A) Representative images of GluN1/GluN3A, nonfunctional GluN1^ΔM4^/GluN3A, and GluN1^ΔM4^+M4^N1^/GluN3A receptor combinations. M4^N1^ failed to rescue functionality in M4-truncated GluN1/GluN3A receptors. (B) SDS-PAGE of metabolic [35S]methionine-tagged GluN1/GluN3A and GluN1^ΔM4^/GluN3A receptors with simultaneous purification of C-terminal His-tagged receptor subunit constructs by metal affinity chromatography (His-tag purification applies to B-E.). Both the GluN1/GluN3A and GluN1^ΔM4^/GluN3A receptors were correctly expressed. (C and D) SDS-PAGE of metabolic [35S]methionine-labeled GluN1/GluN3A receptor combinations. Glycosylation status was determined by PNGaseF or EndoH treatment and showed that GluN1^ΔM4^/GluN3A and GluN1/GluN3A^ΔM4^ were N-glycosylated, indicating cell surface localization. Coexpression of the respective M4 segment resulted in impaired receptor expression or biogenesis in the ER. (E) SDS-PAGE of Cy5-stained surface proteins and GluN1/GluN3A and GluN1^ΔM4^/GluN3A and GluN1^ΔM4^+M4^N1^/GluN3A receptors purified via His-tag with/without PNGase F or EndoH treatment. GluN1/GluN3A and GluN1^ΔM4^/GluN3A were N-glycosylated and presented well at the surface. Coexpression of the M4 segment resulted in complete loss of surface expression. (F) SDS-PAGE was performed with the surface proteins isolated based on biotin affinity purification. (Valid for F-H.). Western blot performed with a primary antibody against the GluN1-NTD shows well surface-expressed GluN1/GluN3A and GluN1^ΔM4^/GluN3A receptors and a complete loss of surface expression for M4 segment co-expressed GluN1^ΔM4^+M4^N1^/GluN3A. (G) Western blot performed with a primary antibody against the GluN1-NTD shows correct surface expression for GluN1/GluN2A, GluN1^ΔM4^/GluN2A, and GluN1^ΔM4^+M4^N1^/GluN2A without affecting M4 segment co-expression. (H) Western blot analysis using a primary antibody against the GluN1 CTD showing that for GluN1^ΔM4^+M4^N1^/GluN2A and M4^N1^ expression alone, the M4 segment was well expressed at the cell surface. The GluN1/GluN2^ΔM4^ was also well expressed at the surface.

Interestingly, however, we noticed a differential shift in the molecular masses of the GluN1^ΔM4^/GluN3A and GluN1/GluN3A^ΔM4^ receptor proteins upon treatment with PNGase F and EndoH (see Figs. 2C and D, lanes 2 and 3), indicating surface expression of the receptors in the absence of M4. We therefore performed surface labeling experiments with affinity purification of GluN1^ΔM4^/GluN3A-His receptors from Cy5 surface-labeled oocytes using a Cy5-NHS ester-based protocol (Schüler et al., 2008). Affinity purification of GluN1^ΔM4^/GluN3A receptors confirmed our assumption that already the GluN1^ΔM4^ and GluN3A subunits, just like GluN1/GluN3A receptors are efficiently located at the cell surface (Fig. 2E, lanes 5, 6 and lanes 8, 9). Consistent with our results with the [35S]methionine-labeled subunits (see Fig. 2C), analysis of the surface-labeled GluN1ΔM4/GluN3A receptors showed a complete loss of surface expression for GluN1^ΔM4^/GluN3A receptors after coexpression with the M4^N1^ fragment (Fig. 2E, lanes 1-3). We therefore planned further surface labeling experiments to investigate whether the difference in rescue of GluN1^ΔM4^/GluN3A and GluN1^ΔM4^/GluN2A channel function in the presence of the M4^N1^ fragment could be explained by a loss of plasma membrane insertion of the GluN1^ΔM4^/GluN3A surface receptors. Thus, we performed cell surface biotinylation of GluN1^ΔM4^/GluN3A- and GluN1^ΔM4^/GluN2A receptor-expressing oocytes, followed by separation of the nonbiotinylated intracellular proteins using a streptavidin agarose pull-down approach. Western blot analysis of the bound biotinylated protein fraction with a primary antibody against the GluN1 extracellular epitope revealed specific bands for GluN1 (125 kDa) and GluN1^ΔM4^ (110 kDa) subunits after expression with the GluN3A and GluN2A subunits, respectively (Fig. 2F, G). Interestingly, GluN1^ΔM4^ protein was absent at the surface when expressed in GluN1^ΔM4^/GluN3A receptors in the presence of the M4^N1^ fragment (Fig. 2F), whereas surface-tagged GluN1^ΔM4^ protein was found when coexpressed in GluN1^ΔM4^/GluN2A receptors with the M4^N1^ fragment (Fig. 2G). Metabolic labeling of a GluN2A*-His construct (see Material and Methods) and the GluN1^ΔM4^ subunit in the absence and presence of M4^N1^ showed no difference in the [35S]methionine-labeled subunit bands and no remarkable decrease in total protein concentrations (see Suppl. Material). This is in marked contrast to our results obtained with the expressed GluN1^ΔM4^/GluN3A subunits, where expression was abolished in the presence of M4^N1^ (Fig. 2C). Apparently, unlike the GluN1^ΔM4^/GluN3A receptors, the coexpressed M4^N1^ fragment does not interfere with the surface expression/assembly of the GluN1^ΔM4^/GluN2A receptor complex. We therefore hypothesized that GluN1^ΔM4^/GluN2A receptors are converted in the plasma membrane into functional receptors by interaction with M4^N1^ (see Fig. 1B). Therefore, we performed cell surface biotinylation of GluN1^ΔM4^/GluN2A receptor-expressing oocytes with M4^N1^ followed by Western blot analysis of the bound biotinylated protein fraction using a primary antibody against the C-terminal (CTD) epitope of GluN1. By using this specific CTD antibody, we detected a protein band of 15 kDa representing our M4^N1^ fragment (Fig. 2H). Interestingly, surface expression of this M4^N1^ protein was indeed seen not only in GluN1^ΔM4^/GluN2A-expressing oocytes but also upon single-cell expression (Fig. 2H), suggesting that our M4^N1^ construct is efficiently transported to the cell surface in the absence of the core receptor and can interact directly with the nuclear receptor in the cell membrane, which could render M4-deleted GluN1^ΔM4^/GluN2A receptors functional.

Thus, our analyses of surface-labeled M4-deleted NMDA receptors demonstrate that both M4-deleted glutamate-gated GluN1/GluN2A and glycine-gated GluN1/GluN3A receptors, although nonfunctional, are efficiently localized to the cell surface. The restoration of channel function for GluN1^ΔM4^/GluN2A- or GluN1/GluN2A^ΔM4^ receptors by selected M4 fragments and the surface localization of singly expressed M4 domains further suggest that our M4 constructs may lead to functional channels through interaction with surface receptors. This does not appear to be true for rescue of channel function in GluN1/GluN3A receptors lacking the M4 domain, because expression of the respective M4 leads to loss of expression of M4-truncated GluN1/GluN3A receptors. In summary, we suggest that M4-deleted GluN1/GluN2A receptors are correctly aligned at the plasma membrane but are nonfunctional, and that coexpression of the corresponding M4 could rescue channel function by interacting with the core receptor in the plasma membrane.

### 3.3 Mapping of functional M4 interfaces of the GluN1 subunit in GluN1/GluN2A receptors

Taken together, our previous data underscore the specificity of M4 transmembrane segment interactions for restoring receptor function rather than receptor assembly. Because coexpression of the M4^N3A^ fragment, in contrast to the M4^N1^ and M4^N2A^ fragments, failed to restore M4-deleted GluN1/GluN2A NMDA receptor function, we interpreted this result as strong evidence that specific interactions with the M4 domain are present, mediated by residues that are not conserved in the GluN3-M4 domain. We therefore analyzed a single alanine substitution at the methionine 813 position in M4^N1^, which is highly conserved in the GluN1 and GluN2 subunits but is a phenylalanine in GluN3 (Fig. 3A). Coexpression of the M4^N1-M813A^ fragment with GluN1^ΔM4^/GluN2A receptors showed little rescue effect compared with wt M4^N1^ (Fig. 3B). Substitution of phenylalanine 817 (conserved in all NMDAR subunits; Fig. 3A) by alanine in M4^N1-F817A^ still resulted in reduced rescue of channel function when coexpressed with the GluN1^ΔM4^/GluN2A receptor, whereas substitution of methionine 818 (tyrosine in GluN2 and valine in GluN3) in M4^N1-M818A^ and leucine 819 (methionine in GluN2) in M4^N1-L819A^ showed a similar rescue effect as M4^N1^ (Fig. 3E). To exclude the possibility that differences in surface expression of the M4^N1^, M4^N1-M813A^, and M4^N1-F817A^ fragments were the cause of the differential rescue of GluN1^ΔM4^/GluN2A channel function, we performed cell surface biotinylation followed by Western blot analysis. For all three M4^N1^ constructs, we detected protein bands of 15 kDa with similar intensities (Fig. 3C). This shows that all of our M4^N1^ mutants are efficiently expressed and transported to the cell surface, suggesting that substitution of methionine 813 and phenylalanine 817 specifically affects M4 interaction with the core receptor. Based on the crystal structures of GluN1/GluN2 NMDA receptors, the ability of our M4^N1^ fragments to rescue the channel function of M4-shortened GluN1^ΔM4^/GluN2A NMDA receptors is consistent with a highly specific interaction between M4^N1-M813^ and the transmembrane segments of each adjacent GluN2A subunit. To identify the molecular basis of M4^N1-M813^ interactions in GluN1/GluN2A receptors, we mutated residues in the TMs of GluN2A that were identified as possible candidates based on i) an analysis of published structures of the transmembrane domains of GluN1/GluN2 receptors (Lü et al., 2017) and ii) a sequence alignment of the TMs of GluN2 NMDA receptor subunits (Fig. 3A). Visual inspection of the transmembrane interactions of methionine 813 of GluN1 with residues of the TMs of the GluN2A subunit based on the crystal structure of the transmembrane domains of the tetrameric GluN1/GluN2A/GluN2B receptor (entry 5UP2 in the Protein Data Bank) revealed, that phenylalanine 637 in M3 of GluN2A (conserved in all GluN2 subunits), is located in the M4/M3 interface of adjacent GluN1 and GluN2 subunits with a Cα-distance in the range of 6 Å (Fig. 3D). Coexpression of M4^N1^ in GluN1^ΔM4^/GluN2A^F637A^ receptors resulted in impaired functional rescue (Fig. 3E). In contrast, coexpression with GluN1^ΔM4^/GluN2A^F641A^ receptors with a mutation of phenylalanine 641 in M3, which is not located near M813 according to structural analysis, resulted in complete functional rescue (Fig. 3E), indicating a specific GluN1 M4^-M813^-GluN2A M3^-F637^ interaction.

**Figure 3:**
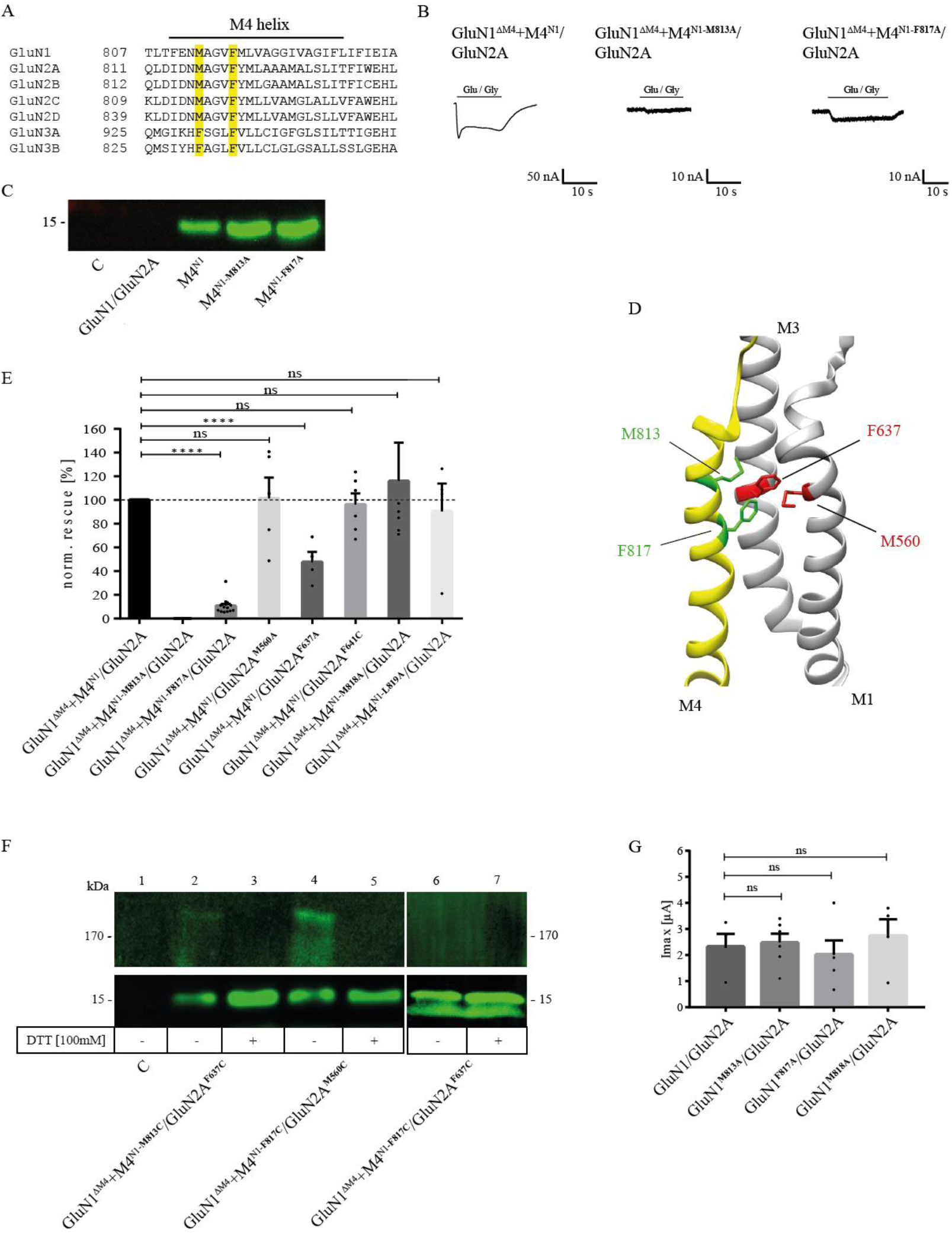
Examination of the possible interaction residues of GluN1-M4 and GluN2A-M1/M3. (A) Sequence alignment of the human GluN1, GluN2 and GluN3 M4 domains. Sequences were taken from Uniprot and sequence alignment was performed using the Multiple Sequence Alignment Tool from EMBL-EBI. The residues identified as possible attachment sites, M813 and F817, are highlighted in yellow. (B) Representative TEVC images of GluN1^ΔM4^+M4^N1^/GluN2A, GluN1^ΔM4^+M4^N1-M813A^/GluN2A, and GluN1^ΔM4^+M4^N1-F817A^/GluN2A show the impaired rescue effect of the two point mutations. M4^N1-M813A^ resulted in a complete loss of rescue effect, while M4^N1-F817A^ resulted in a ~90% decrease in rescued current. (C) Western blot of surface proteins isolated by biotinylation, with a primary antibody against the GluN1 CTD, showing that M4^N1^, M4^N1-M813A^, and M4^N1-F817A^ are all well expressed at the cell surface, ruling out that impaired expression is responsible for the loss of the rescue effect. (D) Representation of GluN1-M4 (yellow) and GluN2A-M1/M3 (gray). Structural analysis was performed using UCSF Chimera with the GluN1/GluN2A structure 5UP2. Amino acids M813 and F817 are colored green, and potential interaction partners M1-M560 (partner of F817) and M3-F637 (partner of M813) are highlighted in red. (E) Quantification of the rescue effect of alanine substitutions normalized to the unmutated M4 segment in co-expressed M4^N1^ showed that for M4 mutations M818A and L819A, in contrast to M4^N1-M813A^ and M4^N1-F817A^, there was no effect on the rescued currents. The GluN2A^M560A^ mutation also showed no effect on the rescue effect, suggesting a stronger influence of the F817-M560 interaction. GluN2A^F637A^ showed a strong decrease in rescued Imax currents by ~50 %, highlighting the importance of the M813-F637 interaction for M4 binding to the truncated nuclear receptor. Statistics by unpaired t-test. (F) Western blot of isolated surface proteins, with a primary antibody against the GluN1-CTD. Samples were equally divided and loaded untreated and DTT-treated. The potential interaction partners M813 - F637 and F817 - M560 were mutated to cysteines to confirm close proximity in the GluN1/GluN2A receptor. GluN1^ΔM4^+M4^N1-M813C^/GluN2A^F637C^ and GluN1^ΔM4^+M4^N1-F817C^/GluN2A^M560C^ both showed both a ~180 kDa and a ~15 kDa band under denaturing conditions without DTT. In both cases, the 180 kDa band disappeared after DTT treatment with an increase in the 15 kDa band, indicating a disulfide bond between the M4^M813C^ (or F817C) with the GluN2A^F637C^ (or M560C). The GluN1^ΔM4^+M4^N1-F817C^/GluN2A^F637C^ sample loaded as a control did not result in a band around the 180 kDa mark, indicating that no disulfide bond was formed here. (G) Quantification of I_max_ currents of wt and alanine-substituted full-length GluN1/GluN2A receptors showing no significant decrease, indicating that the M813A, F817A, and M818A mutations do not affect receptor activity in the full-length receptor. Statistics by unpaired t-test. Data represent mean ±SEM values.

To test whether in GluN1/GluN2A receptors the GluN1-M813 position is indeed close to the amino acid GluN2A-F637, we replaced these residues with cysteines and generated M4^N1-M813C^ and GluN2A^F637C^ constructs. To analyze the M4^N1-M813C^-GluN2A^F637C^ interactions, we performed cell surface biotinylation of GluN1^ΔM4^/GluN2A^F637C^ with M4^N1-M813C^-expressing oocytes and Western blot analysis with the primary antibody against the C-terminal epitope of GluN1 under denaturing conditions in the absence of dithiothreitol (DTT). This revealed both a ~ 180- and a 15-kDa band, indicating the disulfide-linked M4^N1-M813C^ subunit GluN2A^F637C^ and individual M4^N1-M813C^ fragments expressed at the cell surface, respectively (Fig. 3F). The 180-kDa band decreased in the presence of DTT, whereas the intensity of the 15-kDa band was significantly increased (Fig. 3F). This suggests that in GluN1^ΔM4^+M4^N1-M813C^/GluN2A^F637C^ receptors, the introduced cysteines form a disulfide bond that can be released in the presence of DTT, resulting in the loss of the 180-kDa band of the disulfide-bonded M4^N1-M813C^ fragment to GluN2A^F637C^. Because the M4^N1-F817A^ fragment also showed reduced recovery of functionality when coexpressed with GluN1^ΔM4^/GluN2A receptors and is also localized at the TMD interface of GluN1/GluN2A receptors (see Protein Data Bank crystal structure entry 5UP2), we also treated oocytes with DTT expressing the M4^N1-F817C^ and GluN2A^M560C^ mutant, a residue located at the M4/M1 interface of the GluN1 and GluN2A subunits, with a Cα-distance of 6Å from F817 (Fig. 3D). Similar to the GluN1^ΔM4^+M4^N1-M813C^/GluN2A^F637C^ receptors, a ~180 kDa band decreased in the presence of DTT, whereas the intensity of the 15 kDa band increased again (Fig. 3F). In contrast, coexpression of M4^N1-F817C^ with GluN1^ΔM4^/GluN2A^F637C^ did not result in disulfide bond formation (Figs. 3F lanes 6, 7). Thus, the GluN1-M813/GluN2A-F637 and GluN1-F817/GluN2A-M560 positions are capable of forming a specific disulfide bond. However, coexpression of both GluN1^ΔM4^/GluN2A^F637C^ with M4^N1-M813C^ and GluN1^ΔM4^/GluN2A^M560C^ with M4^N1-F817C^ showed no current responses to saturating concentrations of agonists in both the absence and presence of DTT (n=10) (data not shown). This suggests that the disulfide bonds between transmembrane domains may not be resolved in native cell surface-anchored receptors and that the Cα atoms of positions GluN1-M813 and GluN2A-F637 or GluN1-F817 and GluN2A-M560 in GluN1/GluN2A receptors are separated by only about 6 Å. To gain insight into the functional role of the identified interdomain residues in the GluN1/GluN2A receptor, we mutated residues M813, F817, and M818 in the M4 of GluN1-M4, F637 and F641 in the M3, and M560 in the M1 of GluN2A to alanine to analyze the functional properties of the mutants GluN1^M813A^, GluN1^F817A^, GluN1^M818A^, GluN2A^M560A^, GluN2A^F637A^, and GluN2A^F641A^ after coexpression with the respective wt subunit by TEVC. Interestingly, all mutants showed I_max_ currents and glutamate EC_50_ values comparable to wt-GluN1/GluN2A receptors (mean I_max_ ranging from 2.0 to 2.7μA; Fig. 3G and data not shown). In conclusion, the interface mutants of the M4 of GluN1 with the M1 or M3 of GluN2A identified here possess a remarkable role in the association or interaction of the M4 of GluN1 expressed as an isolated peptide with the adjacent GluN2A subunit, whereas little effect of the point mutations on the function within the complete subunits was detected.

### 3.4 Role of the GluN1-M4 interfaces in the action of pregnenolone sulfate

To investigate the importance of our identified interactions of the M4 domains with the M1 and M3 domains of the neighboring subunit for the functional modulation of GluN1/GluN2A receptors, we performed Imax measurements in the absence and presence of the neurosteroid pregnenolone sulfate (20-oxo-5-pregnen-3β-yl sulfate, abbreviated PS; (Horak et al., 2006; Jang et al., 2004; Krausova et al., 2020; Wilding et al., 2016)) after expression of the wt-GluN1/GluN2A receptor and the GluN1^ΔM4^+M4^N1^/GluN2A and GluN1/GluN2A^ΔM4^+M4^N2A^ constructs at saturating agonist concentrations. Remarkably, when I_max_ values were compared in the absence and presence of PS for the different constructs, a difference in the modulation of whole-cell currents was immediately apparent. In contrast to the GluN1/GluN2A and GluN1/GluN2A^ΔM4^+M4^N2A^ receptors, in which PS at a concentration of 100 μM only changed the maximal agonist-inducible whole-cell currents by a maximum of 1.1-fold (Fig. 4A, B), the I_max_ values of GluN1^ΔM4^+M4^N1^/GluN2A receptors were extremely increased by 2.89±0.20-fold in the presence of PS (t(14) =8.739; p <0.0001; Fig. 4A, B). Analysis of M4^N1^ mutants M813A, F817A, M818A, and L819A in the presence of PS revealed an increase in Imax for M4^N1-F817A^ by 2.67±0.12-fold, for M4^N1-M818A^ by 5.93±1.21-fold, and for M4^N1-L819A^ by 2.60±0.21-fold after coexpression with GluN1^ΔM4^/GluN2A (Fig. 4B). For GluN1^ΔM4^+M4^N1-M813A^/GluN2A, no current could be measured even after PS addition (data not shown), again highlighting the particular importance of the M813 position for binding of the M4 domain to the GluN1^ΔM4^/GluN2A nuclear receptor. To decipher possible differences in the mechanism of PS modulation of GluN1/GluN2A- and GluN1^ΔM4^+M4^N1^/GluN2A-mediated currents, we analyzed glutamate dose-response curves in the presence of potentiating PS concentrations. This revealed that PS induced a similar 2-fold increase in apparent glutamate affinity for both GluN1/GluN2A and GluN1^ΔM4^+M4^N1^/GluN2A receptors (Suppl. Data). Thus, a shift in glutamate EC_50_ value from 4.2±0.47 μM to 1.87±0.29 μM and from 2.3±0.2 μM to 1.31±0.2μM was observed for GluN1/GluN2A and GluN1^ΔM4^+M4^N1^/GluN2A receptors, respectively. Similarly, analysis of PS potentiation affinity for the GluN1/GluN2A and GluN1^ΔM4^+M4^N1^/GluN2A receptors revealed nearly similar EC50 values of 12.4±1.3 μM and 15±1.1 μM, with a small but significant decrease in PS affinity for GluN1^ΔM4^+M4^N1^/GluN2A (Fig. 4C; t(6) =2.873; p =0.0283). Interestingly, the M4^N1^ mutant M818A, which showed the strongest increase in I_max_ of 5.93±1.21-fold in the presence of PS (see Fig. 4B), caused a significant decrease in PS affinity to 23.2±1.4 μM (t(7) =12; p<0.0001). Analysis of GluN1^M813A^/GluN2A and GluN1^M818A^/GluN2A full-length constructs also showed significantly increased Imax potentiation for both compared to wt (1.40±0.09-fold; t(18) =2.461; p=0.0242 and 1.72±019-fold, t(15) =2.99; p=0.0092; Fig. 4D). Since there is evidence for a balance of positive- and negative-modulatory (PAM and NAM) neurosteroid recognition sites leading to PS potentiation in GluN1/GluN2A and GluN1/GluN2B receptors and PS inhibition in GluN1/GluN2C and GluN1/GluN2D receptors (Horak et al., 2006), we examined the effect of PS on the modulation of GluN1/GluN2D and GluN1^ΔM4^+M4^N1^/GluN2D receptors. Significantly, the inhibitory effect of PS at GluN1/GluN2D was converted to a potentiating one at GluN1^ΔM4^+M4^N1^/GluN2D receptors (Figure 4D). These data imply two possibilities; That either i) the NAM effect of PS in GluN1^ΔM4^+M4^N1^/GluN2 receptors is attenuated by a change in the interactions of the M4 of GluN1 with the TMs of neighboring GluN2, and consequently there is enhanced potentiation by PS at the PAM-binding site, or ii) the PAM effect of PS in GluN1^ΔM4^+M4^N1^/GluN2 receptors is enhanced by a change in the interactions of the M4 of GluN1 with the TMs of the neighboring GluN2, resulting in potentiation.

**Figure 4:**
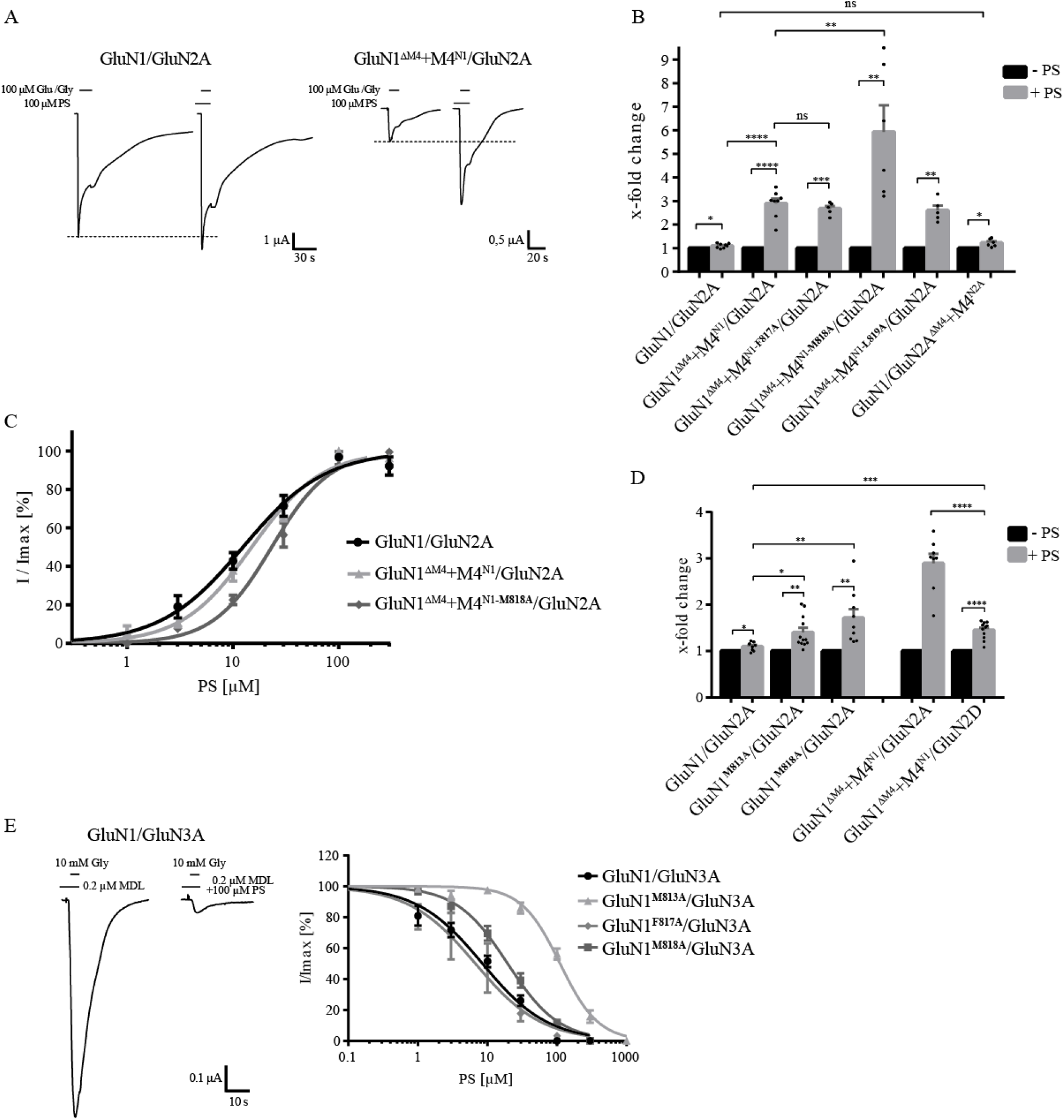
Identification of a NAM recognition site of pregnenolone sulfate at GluN1-M4. (A) Representative TEVC images of I_max_ potentiation of PS at GluN1/GluN2A and GluN1^ΔM4^+M4^N1^/GluN2A receptors showing a strong increase in potentiating effect for M4 segment co-expression. (B) Quantification of Imax potentiation of PS at GluN1/GluN2A, GluN1^ΔM4^+M4^N1^/GluN2A, M4 segment mutants F817A, M818A, and L819A, and GluN1/GluN2A^ΔM4^+M4^N2A^. Coexpression of M4^N1-F817A^ and M4^N1-L819A^ showed no effect on the potentiating effect compared with unmutated M4^N1^. Co-expression of M4^N1-M818A^ showed a strong increase in Imax potentiation compared to GluN1^ΔM4^+M4^N1^/GluN2A (t(12)= 3.08; p= 0.0095), indicating an impaired NAM recognition site. GluN1/GluN2A^ΔM4^+M4^N2A^ was not different from the wt full-length receptor. Statistical analysis was performed using paired t-tests between I_max_ [-PS] and I_max_ [+PS]. Potentiation of the different mutants was analyzed by unpaired t-test. (C) Dose-response analysis of PS for wt GluN1/GluN2A, GluN1^ΔM4^+M4^N1^/GluN2A, and GluN1^ΔM4^+M4^N1-M818A^/GluN2A showing nearly equal EC_50_ values of GluN1/GluN2A and GluN1^ΔM4^+M4^N1^/GluN2A (12.4± 1.3 μM, n = 5 and 15.0± 1.1 μM, n = 3; P<0.05). The M4^N1-M818A^ mutation resulted in a significant decrease in PS affinity (EC_50_: 23.2± 1.4 μM; n = 4; p<0.001) compared to the non-mutated M4^N1^. (D) Quantification of I_max_ potentiation of PS for GluN1/GluN2A, GluN1^M813A^/GluN2A and GluN1^M818A^/GluN2A showed significantly increased potentiation for both mutants compared to wt (M813A 1.40± 0.09-fold; n = 9; p<0.05 and for M818A 1.72± 0.19-fold; n = 12; p<0.01 compared to wt 1.10± 0.09-fold; n = 8). Comparison of GluN1^ΔM4^+M4^N1^/GluN2A and GluN1^ΔM4^+M4^N1^/GluN2D showed a significant difference in PS potentiation (2.89± 0.2-fold, n = 8 compared to 1.45± 0.19-fold, n = 11; p <0.0001). The normally inhibitory overall effect of PS on GluN1/GluN2D was converted to a potentiating one for GluN1^ΔM4^+M4^N1^/GluN2D. (E) Representative TEVC uptake of the inhibition of GluN1/GluN3A by PS. Dose-response curves of PS at GluN1/GluN3A, GluN1^M813A^/GluN3A, GluN1^F817A^/GluN3A, GluN1^M818A^/GluN3A showing no differences between wt and the F817A mutant, but a strong decrease in PS affinity at the M813A and M818A mutants (111± 11 μM, n = 4; p<0. 0001 and 19.5± 1.7 μM; n = 4 p<0.0001 compared to 8.5± 0.9 μM n = 7 for wt). Data represent mean values ±SEM.

To obtain evidence for a possible NAM- or PAM-binding site for PS in the M4 region of GluN1, we examined the effect of PS at glycine-gated GluN1/GluN3 receptors. Thus, in the absence and presence of PS, we performed I_max_ measurements after expression of the wt GluN1/GluN3A receptor following application of the agonist glycine (10 mM) in combination with the potentiating ligand MDL-29951 (0.2 μM). Remarkably, when comparing I_max_ values in the absence and presence of 100 μM PS for the different constructs, a difference in the modulation of whole cell currents of GluN1/GluN2A and GluN1/GluN3A receptors was immediately apparent. In contrast to GluN1/GluN2A receptors, where PS at a concentration of 100 μM potentiated maximal agonist-inducible whole-cell currents (Fig. 4A), I_max_ values of GluN1/GluN3A receptors were extremely decreased in the presence of PS (Fig. 4E). Affinity analysis of PS inhibition for GluN1/GluN3A receptors revealed an IC50 value of 8.5±0.9 μM (n=7), remarkably a similar value to the EC_50_ value of 12.4±1.3 μM for GluN1/GluN2A receptors. Interestingly, GluN1^M813A^ and GluN1^M818A^ mutants showed a strong increase in the IC50 value of PS to 111±11 μM and 19.5±1.7 μM, respectively (t(9) =25.43; p<0.0001 and t(13) =15.22; p<0.0001; Fig. 4E;). In contrast, expression of GluN1^F817A^/GluN3A receptors resulted in an unchanged IC50 value for PS of 6.73±1.73 μM (t(8)=2.178; p=0.061). Thus, our analysis of GluN1^M813A^/GluN3A and GluN1^M818A^/GluN3A receptors revealed strong evidence for a NAM binding site for PS at the interface of the M4 of the GluN1 subunit with the adjacent GluN3A subunit. Considering the assumption of a PAM and NAM neurosteroid recognition site in GluN1/GluN2 receptors, the increase in PS potentiation of our GluN1-M4 mutations in GluN1/GluN2A receptors implies an impairment of inhibition by the NAM-PS binding-site at the interface of the M4 of the GluN1 subunit to the neighboring GluN2A subunit. Consequently, impairment of inhibition by PS leads to an increase in potentiation by PS at the corresponding PAM-binding site. Thus, our results suggest a specific role of amino acid residues in GluN1-M4 for the binding of the M4 domain to the nuclear receptor, which are involved both in stabilizing the interactions of the isolated M4 to the nuclear receptor and in the negative modulatory effect of the neurosteroid PS.

## 4. Discussion

Due to partially contradictory results, the importance of the transmembrane domain M4 in ionotropic glutamate receptor function and assembly remains controversial. In the present study of the role of M4 in NMDA receptor functionality, we show that deletion of M4 in the GluN1, GluN2A, and GluN3A subunits, despite retained receptor assembly and surface expression, results in nonfunctional membrane receptors. Remarkably, coexpression of the corresponding M4 domains of GluN1 and GluN2A, but not of the GluN3A subunit, as an isolated peptide in M4-deleted GluN1/GluN2A receptors rescued receptor function without altering agonist affinities. The substitution of non-conserved residues within the putative interfaces of M4 of GluN1 with neighboring GluN subunits suggest a specific role of these residues in i) the coupling of isolated-expressed M4 fragments to the core receptor and ii) the negative modulatory effect of the neurosteroid pregnenolone sulfate, highlighting the importance of M4 and its interactions in the regulation of NMDA receptor function.

### 4.1 The role of M4 in NMDAR assembly and function

Although there are numerous reports on the involvement of M4 in the assembly of iGluRs, its role remains controversial. In all iGluRs, the M4 domain of one subunit is structurally linked to the pore-forming M1 and M3 helices of the neighboring subunit (Karakas and Furukawa, 2014; Sobolevsky et al., 2009). Based on this exclusive interaction of the peripheral M4 domain with the central M1 and M3 of the neighboring subunit, a strong interaction of residues of the M4 domain with these transmembrane domains has been proposed to mediate receptor assembly or at least surface targeting of iGluRs (Amin et al., 2017). Indeed, some results showed that the peripheral M4 helix is involved in subunit association or at least confers additional stability to the tetrameric receptor, ideas based mainly on AMPA receptor assembly studies (Amin et al., 2017; Gan et al., 2016; Salussolia et al., 2011, 2013). In contrast, studies on conventional GluN1/GluN2 NMDA receptors led to the view that the M4 domain is more required for the formation of GluN1/GluN2 heterodimers and, in the case of the GluN2B subunit, also for masking ER retention signals in GluN1 (Cao et al., 2011; Horak et al., 2008). We can show that no M4 deletion affects the assembly and surface expression of the conventional GluN1/GluN2A NMDA receptor, supporting the view that the M4 of glutamate-gated NMDARs are primarily structural determinants that are more involved in the allosteric regulation of ion channel opening by modulatory compounds (Krausova et al., 2020). The absence of M4 in prokaryotic GluR0 also supports the idea that this transmembrane region may not be essential for iGluR assembly and that its presence in eukaryotic iGluRs is predominantly required for modulating or fine-tuning the kinetic properties of the channel. Consistent with this conclusion, our current study also shows for the less studied glycine-gated GluN1/GluN3A NMDA receptor that deletion of M4 does not affect assembly or cell surface expression. In summary, our data strengthen the view that the M4 domain is not required for oligomerization of glutamate- or glycine-gated NMDA receptors.

Thus, although we found that the M4 domain is not involved in NMDA receptor assembly and surface trafficking, it is essential for receptor function. This is impressively demonstrated by the rescue of channel function of M4-deleted GluN1/GluN2A receptors with unaltered agonist affinity after coexpression with the corresponding M4 domain as an isolated peptide. Surprisingly, coexpression of M4 in GluN1/GluN3A M4-deleted constructs resulted in a complete loss of receptor subunit expression, although all M4 segments tested were expressed and incorporated into the cell surface with equal efficiency. We interpret this pronounced instability of M4-deleted GluN1/GluN3A receptor proteins in the presence of the corresponding M4 domain as indicating rapid degradation of these receptors in the ER by quality control mechanisms. In contrast, for the M4-deleted GluN1/GluN2A receptors, we found retained surface expression after coexpression with the corresponding M4 domain, which, however, is accompanied by a decrease in maximal inducible whole cell currents, a finding already described in a previous study on M4-deleted NMDARs (Schorge and Colquhoun, 2003). This may be due to a reduced likelihood of selective interaction of surface-expressed nuclear receptors and isolated M4 fragments. This is also supported by our finding that coexpression of the M4 of GluN1 could also rescue GluN1/GluN2A^ΔM4^ receptors, whereas the less conserved M4 segment of the GluN3A subunit in GluN1^ΔM4^/GluN2A or GluN1/GluN2A^ΔM4^ could not restore receptor function. This ability of M4 segments to differentially rescue the functionality of M4-deleted GluN1/GluN2 receptors could be determined by the individual exchange of amino acid residues in the interface of M4 with neighboring TMs. Thus, exchange of residue M813 conserved in the GluN1 and GluN2 subunits in M4 of GluN1, a methionine associated with refractory seizures and global developmental delay, when exchanged for valine in the GluN2A subunit (Chen et al., 2017; Venkateswaran et al., 2014), specifically decreased Imax upon coexpression of M4-deleted receptors but not in full-length mutant GluN1/GluN2A receptors, again suggesting a specific role of interactions in the likelihood of functional coupling of M4 to the nuclear receptor. This is also supported by our finding that the nearby residue F637 in M3 of the neighboring GluN2 subunit also decreased the I_max_ of M4-deleted receptors. Consistent with our findings, the insertion of large residues in a 2017 study by Amin and colleagues in a tryptophan scan of the M4 interface of the GluN1 and GluN2A subunits also confirms decreased functionality of the M4 interfaces.

Overall, we identified pairs of amino acid residues important for the interactions of M4 with the core receptor and involved in the restoration of receptor function by isolated M4 peptides. We confirmed phenylalanine 637 of M3 of the GluN2A subunit as an interaction partner of methionine 813 by forming a disulfide bond after cysteine substitution (GluN1-M813C and GluN2A-F637C). The second close interaction is between GluN1-F817 of M4 and position M560 at M1 of the GluN2A subunit. GluN1^ΔM4^+M4^N1-F817A^/GluN2A showed a strong reduction in rescue effect (~90 %), although not a complete loss of function as with M4^N1-M813A^. Moreover, GluN1-F817 with a phenylalanine to leucine exchange is known to be disease-associated, leading to intellectual and mental disability, highlighting its importance for NMDAR functionality (Lemke et al., 2016). Cysteine substitution at the two residues (M4^N1-F817C^ and GluN2A^M560C^) again allowed insertion of a disulfide bond. Steric restriction of the TM interface by oxidative cross-linking in the GluN1^ΔM4^+M4^N1-M813C^/GluN2A^F637C^ and GluN1^ΔM4^+M4^N1-F817C^/GluN2A^M560C^ receptors resulted in loss of function in both, suggesting that the receptor interface requires some degree of flexibility. Our assumptions are consistent with i) X-ray crystal structure analyses of GluN1/GluN2 receptor complexes showing a unique arrangement of M4 domains with distinct intersubunit interactions (Lee et al., 2014) and ii) Wollmuth lab molecular dynamics simulations showing that the top of the M4 helix must move to stabilize the open state of the NMDA receptor (Amin et al., 2018). Based on the results presented here, we propose that the main role of M4 in NMDA receptors is to ensure the functionality of agonist-induced channel opening. Our results clearly show that M4 is not involved in the efficiency of assembly; in contrast, M4 with its interactions rather represents an important domain for conformational changes within TMs for channel opening and their modulation after ligand binding at the interface of TMs.

### 4.2 The role of M4-TM interfaces in determining the efficiency of pregnenolone sulfate (PS) modulation on NMDARs

The importance of interactions at TM interfaces in the binding and modulation of compounds to NMDARs is poorly understood. As mentioned above, M4 interfaces are thought to alter the kinetic properties of conventional NMDARs by rearranging the M1 and M4 helices, thereby stabilizing the open-state position of the M3 helices (Amin et al., 2018, 2017) In addition, NMDAR channel function is likely to be strongly modulated by repositioning of peripheral M4 segments through interactions with lipids or through binding sites for positive and negative modulators (Casado and Ascher, 1998; Korinek et al., 2015; Ren et al., 2012; Traynelis and Wollmuth L.P., McBain C.J., Menniti F.S., Vance K.M., Ogden K.K., Hansen K.B., Yuan H., Myers J.M., 2010; Wilding et al., 2016). We can show that the M813A interface mutation in M4 of GluN1 specifically decreases inhibition of GluN1/GluN2A, and particularly pronounced, at GluN1/GluN3A receptors by PS. Our results further clearly demonstrate that the M4 of GluN1 and the M4 of GluN2 may not be equally involved in determining neurosteroid efficiency; rather, the M4 of GluN1 represents an important domain for the negative effect of PS. Consequently, we attribute the complete lack of a positive-modulatory effect of PS in GluN1/GluN3A receptors to the unique design of the M4 interface of the GluN3 subunit. Our results are consistent with other studies of conventional GluN1/GluN2 NMDA receptors, in which it has been shown that the M4 and its linker region of GluN2 subunits, in particular, control the subunit-specific PS action by determining the positive-modulatory effect of PS (Jang et al., 2004; Krausova et al., 2020). However, the exact mechanism that couples the modulatory properties of the M4 domain to channel activation and the importance of these TM interfaces in subunit-specific PS regulation are still unknown. In the absence of detailed information on the structure and conformation of the M4 interface upon binding of a modulator within TMs, we hypothesize that specific side-chain interactions within the M4/TM regions are important for positive-or negative-regulatory interactions. We think that our proposed negative-modulatory steroid interaction site, formed by the M4-GluN1 and M1/M3-GluN2 or GluN3 helices, undergoes a structural rearrangement after binding of PS and thus negatively-allosterically affects channel conformation. Ultimately, this negative-modulatory effect would reduce the potentiating effect of a second, distinct steroid-binding site. Based on the results presented here, we therefore propose that i) the flexibility and positioning of the M4 of GluN1 is important for the inhibitory effect of PS, and ii) that the ratio of the effects of the positive- and negative-modulatory steroid-binding site determines the subunit-dependent modulation of GluN1/GluN2/3 receptors by PS.

### Conclusions

The present study demonstrates a prominent role of the M4 of GluN1 in determining PS efficacy at NMDA receptors, because mutations within the TM interfaces result in a strong loss of PS-induced inhibition in GluN1/GluN3A receptors. Taken together, our results implicate distinct roles of M4 domains in different NMDA receptor subunits and highlight their importance in the regulation of NMDA receptor function by neurosteroids. Compounds with PS-like properties targeting the GluN1-M4 interfaces may represent powerful tools for selective modulation of glutamate- and glycine-activated NMDA receptors *in vivo*.

## ACKNOWLEDGMENTS

This study has been supported in the frame of the LOEWE project iNAPO by the Hessen State Ministry of Higher Education, Research and the Arts. The authors are grateful to Michael Schönrock for critical reading of the manuscript. The authors acknowledge support by the German Research Foundation and the Open Access Publishing Fund of Technische Universität Darmstadt.

## AUTHORS CONTRIBUTIONS

KL, AML, and JW designed and performed experiments; KL, AML, JW and BL analyzed data; KL and BL prepared the manuscript. All authors approved the final version of the manuscript.

## REFERENCES

Amin, J.B., Leng, X., Gochman, A., Zhou, H.X., Wollmuth, L.P., 2018. A conserved glycine harboring disease-associated mutations permits NMDA receptor slow deactivation and high Ca2+ permeability. Nat. Commun. 9. https://doi.org/10.1038/s41467-018-06145-w

Amin, J.B., Salussolia, C.L., Chan, K., Regan, M.C., Dai, J., Zhou, H.X., Furukawa, H., Bowen, M.E., Wollmuth, L.P., 2017. Divergent roles of a peripheral transmembrane segment in AMPA and NMDA receptors 1–20. https://doi.org/10.1085/jgp.201711762

Cao, J.Y., Qiu, S., Zhang, J., Wang, J.J., Zhang, X.M., Luo, J.H., 2011. Transmembrane region of N-Methyl-D-aspartate receptor (NMDAR) subunit is required for receptor subunit assembly. J. Biol. Chem. 286, 27698–27705. https://doi.org/10.1074/jbc.M111.235333

Casado, M., Ascher, P., 1998. Opposite modulation of NMDA receptors by lysophospholipids and arachidonic acid: Common features with mechanosensitivity. J. Physiol. 513, 317–330. https://doi.org/10.1111/j.1469-7793.1998.317bb.x

Chen, W., Tankovic, A., Burger, P.B., Kusumoto, H., Traynelis, S.F., Yuan, H., 2017. Functional evaluation of a de novo GRIN2A mutation identified in a patient with profound global developmental delay and refractory epilepsy. Mol. Pharmacol. 91, 317–330. https://doi.org/10.1124/mol.116.106781

Collingridge, G.L., Bliss, T.V.P., 1995. Memories of NMDA receptors and LTP. Trends Neurosci. 18, 54–56. https://doi.org/10.1016/0166-2236(95)80016-U

Gan, Q., Dai, J., Zhou, H.X., Wollmuth, L.P., 2016. The transmembrane domain mediates tetramerization of α-Amino-3-hydroxy-5-methyl-4-isoxazolepropionic Acid (AMPA) Receptors. J. Biol. Chem. 291, 6595–6606. https://doi.org/10.1074/jbc.M115.686246

Honer, M., Benke, D., Laube, B., Kuhse, J., Heckendorn, R., Allgeier, H., Angst, C., Monyer, H., Seeburg, P.H., Betz, H., Mohler, H., 1998. Differentiation of glycine antagonist sites of N-methyl-D-aspartate receptor subtypes: Preferential interaction of CGP 61594 with NR1/2B receptors. J. Biol. Chem. 273, 11158–11163. https://doi.org/10.1074/jbc.273.18.11158

Horak, M., Chang, K., Wenthold, R.J., 2008. Masking of the Endoplasmic Reticulum Retention Signals during Assembly of the NMDA Receptor 28, 3500–3509. https://doi.org/10.1523/JNEUROSCI.5239-07.2008

Horak, M., Vlcek, K., Chodounska, H., Vyklicky, L., 2006. Subtype-dependence of N-methyl-D-aspartate receptor modulation by pregnenolone sulfate. Neuroscience 137, 93–102. https://doi.org/10.1016/j.neuroscience.2005.08.058

Jang, M., Mierke, D.F., Russek, S.J., Farb, D.H., 2004. A steroid modulatory domain on NR2B controls N-methyl-D-aspartate receptor proton sensitivity. PNAS 2–7.

Karakas E and Furukawa H, 2014. Crystal structure of a heterotetrameric NMDA receptor ion channel. Science (80-.). 344, 992–997. https://doi.org/10.1126/science.1251915.Crystal

Korinek, M., Vyklicky, V., Borovska, J., Lichnerova, K., Kaniakova, M., Krausova, B., Krusek, J., Balik, A., Smejkalova, T., Horak, M., Vyklicky, L., 2015. Cholesterol modulates open probability and desensitization of NMDA receptors. J. Physiol. 593, 2279–2293. https://doi.org/10.1113/jphysiol.2014.288209

Krausova, B.H., Kysilov, B., Cerny, J., Vyklicky, V., Smejkalova, T., Ladislav, M., Balik, A., Korinek, M., Chodounska, H., Kudova, E., Vyklicky, L., 2020. Site of action of brain neurosteroid pregnenolone sulfate at the N-methyl-D-aspartate receptor. J. Neurosci. 40, 5922–5936. https://doi.org/10.1523/JNEUROSCI.3010-19.2020

Laube, B., Hirai, H., Sturgess, M., Betz, H., Kuhse, J., 1997. Molecular determinants of agonist discrimination by NMDA receptor subunits: Analysis of the glutamate binding site on the NR2B subunit. Neuron 18, 493–503. https://doi.org/10.1016/S0896-6273(00)81249-0

Lee, C.-H., Wei Lü, Jennifer Carlisle Michel, April Goehring, Juan Du, Xianqiang Song, and E.G., 2014. NMDA receptor structures reveal subunit arrangement and pore architecture 511, 191–197. https://doi.org/10.1038/nature13548.NMDA

Lemke, J.R., Geider, K., Helbig, K.L., Heyne, H.O., Schütz, H., Hentschel, J., Courage, C., Depienne, C., Nava, C., Heron, D., Moller, R.S., Hjalgrim, H., Lal, D., Neubauer, B.A., Nürnberg, P., Thiele, H., Kurlemann, G., Arnold, G.L., Bhambhani, V., Bartholdi, D., Pedurupillay, C.R.J., Misceo, D., Frengen, E., Stromme, P., Dlugos, D.J., Doherty, E.S., Bijlsma, E.K., Ruivenkamp, C.A., Hoffer, M.J. V, Goldstein, A., Rajan, D.S., Narayanan, V., Ramsey, K., Belnap, N., Schrauwen, I., Richholt, R., Koeleman, B.P.C., Sa, J., Mendonca, C., Kovel, C.G.F. De, Weckhuysen, S., Hardies, K., Jonghe, P. De, Meirleir, L. De, Milh, M., Badens, C., Lebrun, M., Busa, T., Francannet, C., Piton, A., Riesch, E., Biskup, S., Vogt, H., Dorn, T., Helbig, I., Michaud, J.L., Laube, B., Syrbe, S., 2016. Delineating the GRIN1 phenotypic spectrum: A distinct genetic NMDA receptor encephalopathy. Neurology 86, 2171–2178. https://doi.org/10.1212/WNL.0000000000002740

Lü, W., Du, J., Goehring, A., Gouaux, E., 2017. Cryo-EM structures of the triheteromeric NMDA receptor and its allosteric modulation. Science (80-.). 355, eaal3729. https://doi.org/10.1126/science.aal3729

Lynagh, T., Kunz, A., Laube, B., 2013. Propofol modulation of α1 glycine receptors does not require a structural transition at adjacent subunits that is crucial to agonist-induced activation. ACS Chem. Neurosci. 4, 1469–1478. https://doi.org/10.1021/cn400134p

Lynagh, T., Laube, B., 2014. Opposing effects of the anesthetic propofol at pentameric ligand-gated ion channels mediated by a common site. J. Neurosci. 34, 2155–2159. https://doi.org/10.1523/JNEUROSCI.4307-13.2014

Madry, C., Mesic, I., Bartholomäus, I., Nicke, A., Betz, H., Laube, B., 2007a. Principal role of NR3 subunits in NR1/NR3 excitatory glycine receptor function. Biochem. Biophys. Res. Commun. 354, 102–108. https://doi.org/10.1016/j.bbrc.2006.12.153

Madry, C., Mesic, I., Betz, H., Laube, B., 2007b. The N-terminal domains of both NR1 and NR2 subunits determine allosteric Zn2+ inhibition and glycine affinity of N-methyl-D-aspartate receptors. Mol. Pharmacol. 72, 1535–44. https://doi.org/10.1124/mol.107.040071

Mesic, I., Madry, C., Geider, K., Bernhard, M., Betz, H., Laube, B., 2016. The N-terminal domain of the GluN3A subunit determines the efficacy of glycine-activated NMDA receptors. Neuropharmacology 105, 133–141. https://doi.org/10.1016/j.neuropharm.2016.01.014

Pettersen, E.F., Goddard, T.D., Huang, C.C., Couch, G.S., Greenblatt, D.M., Meng, E.C., Ferrin, T.E., 2004. UCSF Chimera - A visualization system for exploratory research and analysis. J. Comput. Chem. 25, 1605–1612. https://doi.org/10.1002/jcc.20084

Ren, H., Zhao, Y., Dwyer, D.S., Peoples, R.W., 2012. Interactions among positions in the third and fourth membrane-associated domains at the intersubunit interface of the N-Methyl-D-aspartate receptor forming sites of alcohol action. J. Biol. Chem. 287, 27302–27312. https://doi.org/10.1074/jbc.M111.338921

Salussolia, C.L., Corrales, A., Talukder, I., Kazi, R., Akgul, G., Bowen, M., Wollmuth, L.P., 2011. Interaction of the M4 segment with other transmembrane segments is required for surface expression of mammalian α-amino-3-hydroxy-5-methyl-4-isoxazolepropionic acid (AMPA) receptors. J. Biol. Chem. 286, 40205–40218. https://doi.org/10.1074/jbc.M111.268839

Salussolia, C.L., Gan, Q., Kazi, R., Singh, P., Allopenna, J., Furukawa, H., Wollmuth, L.P., 2013. A Eukaryotic Specific Transmembrane Segment is Required for Tetramerization in AMPA Receptors. J. Neurosci. 33, 9840–9845. https://doi.org/10.1523/jneurosci.2626-12.2013

Schönrock, M., Thiel, G., Laube, B., 2019. Coupling of a viral K+-channel with a glutamate-binding-domain highlights the modular design of ionotropic glutamate-receptors. Commun. Biol. 2, 1–10. https://doi.org/10.1038/s42003-019-0320-y

Schorge, S., Colquhoun, D., 2003. Studies of NMDA receptor function and stoichiometry with truncated and tandem subunits. J. Neurosci. 23, 1151–1158. https://doi.org/10.1523/jneurosci.23-04-01151.2003

Schüler, T., Mesic, I., Madry, C., Bartholoma, I., Laube, B., 2008. Formation of NR1/NR2 and NR1/NR3 heterodimers constitutes the initial step in N-methyl-D-aspartate receptor assembly. J. Biol. Chem. 283, 37–46. https://doi.org/10.1074/jbc.M703539200

Sheng, M., Pak, D.T., 1999. Glutamate receptor anchoring proteins and the molecular organization of excitatory synapses. Ann. N. Y. Acad. Sci. 868, 483–493. https://doi.org/10.1111/j.1749-6632.1999.tb11317.x

Sobolevsky AI., Rosconi MP., and G.E., 2009. X-ray structure of AMPA-subtype glutamate receptor: symmetry and mechanism. Nature 462, 745–756. https://doi.org/10.1038/nature08624.X-ray

Traynelis, S.F., Wollmuth L.P., McBain C.J., Menniti F.S., Vance K.M., Ogden K.K., Hansen K.B., Yuan H., Myers J.M., and D.R., 2010. Glutamate Receptor Ion Channels: Structure, Regulation, and Function. Pharmacol. Rev. 62, 405–496. https://doi.org/10.1124/pr.109.002451.

Venkateswaran, S., Myers, K.A., Smith, A.C., Beaulieu, C.L., Schwartzentruber, J.A., Majewski, J., Bulman, D., Boycott, K.M., Dyment, D.A., 2014. Whole-exome sequencing in an individual with severe global developmental delay and intractable epilepsy identifies a novel, de novo GRIN2A mutation. Epilepsia 55, 75–79. https://doi.org/10.1111/epi.12663

Wilding, T.J., Lopez, M.N., Huettner, J.E., 2016. Chimeric Glutamate Receptor Subunits Reveal the Transmembrane Domain Is Sufficient for NMDA Receptor Pore Properties but Some Positive Allosteric Modulators Require Additional Domains. J. Neurosci. 36, 8815–8825. https://doi.org/10.1523/JNEUROSCI.0345-16.2016

Wo, Z., Oswald, R.E., 1995. Unraveling the modular design of glutamate-gated ion channels. Trends Neurosci. 18, 161–168. https://doi.org/10.1016/0166-2236(95)93895-5

Yashiro, K., Philpot, B.D., 2008. Regulation of NMDA receptor subunit expression and its implications for LTD, LTP, and metaplasticity. Neuropharmacology 55, 1081–1094. https://doi.org/10.1016/j.neuropharm.2008.07.046

