## Supplemental Figures for "Analysis of M4 transmembrane domains in NMDA receptor function: a negative allosteric modulation site at the GluN1 M4 is determining the efficiency of neurosteroid modulation"

**Supplement**

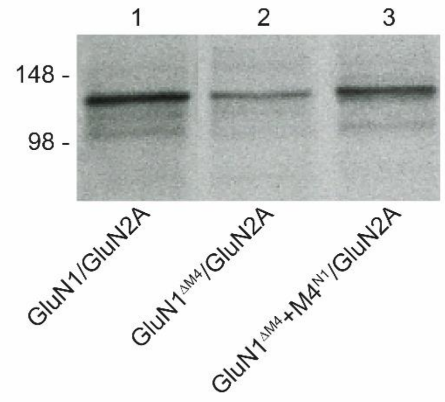


Fig. 1 Impact of M4-truncation and M4-segement coexpression on GluN1/GluN2A receptor expression.

SDS-PAGE of metabolic [35S]methionine-tagged GluN1/GluN2A, GluN1$^{\Delta M4}$/GluN2A and GluN1$^{\Delta M4}$+M4$^{N1}$/GluN2A receptors with purification of C-terminal His-tagged receptor subunit constructs by metal affinity chromatography. All constructs were properly expressed, M4-Segement coexpression did not alter GluN1$^{\Delta M4}$+M4$^{N1}$/GluN2A expression level.

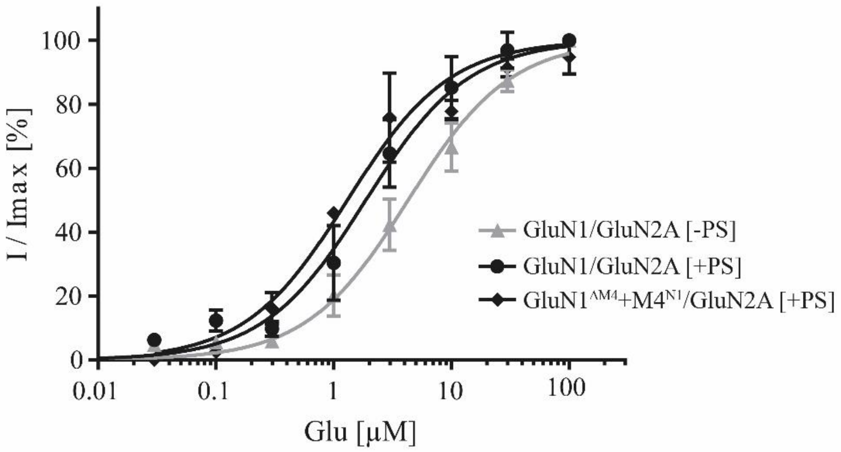


Fig.2 Impact of PS on the Glutamate Affinity.

Dose-response analysis showed that agonist affinity of GluN1/GluN2A and $GluN1^{\Delta M4}+M4^{N1}/GluN2A$ were similar without PS modulation (see results Fig. 1C). PS modulation of both GluN1/GluN2A and $GluN1^{\Delta M4}+M4^{N1}/GluN2A$ resulted in an increase of agonist affinity (GluN1/GluN2A [-PS] $EC_{50}$: 4.2±0.47 µM to [+PS] $EC_{50}$: 1.87±0.29 µM ($t(5) = 7.731$; $p = 0.0006$; and $GluN1^{\Delta M4}+M4^{N1}/GluN2A$ [+PS] $EC_{50}$: 1.31±0.2 µM; $t(6) = 11.64$; $p < 0.0001$). The results show a similar shift of the agonist affinity for both wt and M4-segment coexpression. Statistics done by unpaired t-test. Data represent mean ±SEM.
